# Bacterial Peptidoglycan as a Food Digestive Signal in the Nematode that Facilitates Adaptation of Animals in Nature

**DOI:** 10.1101/2023.03.20.533399

**Authors:** Fanrui Hao, Huimin Liu, Bin Qi

**Author notes:** Corresponding Author (Bin Qi,).

## Abstract

Food availability and usage is a major adaptive force for the successful survival of animals in nature. However, very little is known about the signal from food to activate the hosts digestive system, which facilitates animals to digest more diverse food in nature. Here, by using a food digestion system in *C. elegans*, we discover that bacterial peptidoglycan (PGN) is a unique food signal that activates animals to digest inedible food. We find that PGN was sensed by a conserved intestinal glycosylated protein (BCF-1) in nematodes via direct interaction, which promoted food digestion through inhibiting the mitochondrial unfolded protein response (UPR^mt^). Moreover, constitutive activation of UPR^mt^ is sufficient to inhibit food digestion. Thus, our study reveals how bacterial PGN, as a common digestion cue, activates the food digestive system through interacting with a conserved glycosylated protein, which facilitates adaptation of the host animals by increasing ability to consume a wide range of foods in their natural environment.

## Introduction

Nematodes, including bacterial-feeding nematodes, represent approximately 80% of all the multicellular animals on earth, and they have major roles in many ecosystems (Eisenhauer and Guerra, 2019). In nature, nematode species play an important role in decomposition of soil organic matter, soil aeration, mineralization of plant nutrients, and nutrient cycling (Freckman, 1988). One explanation for their success is their extraordinary ability to consume a wide range of foods.

Food is essential for animal/nematode survival and for regulating their population in nature. Evolutionarily, food availability or preference has been a major adaptive force (Ackley, 2019). Animals have developed a nervous system to sense food for guiding food intake (Davis et al., 2017; Morton et al., 2014; Rhoades et al., 2019). The food is then digested in the gastrointestinal tract to obtain essential nutrients for the animal (Hartenstein and Martinez, 2019). Therefore, a strong digestive system is important for animals to consume almost every available food source, which improves the survival of the animals under difficult circumstances by increasing their ability to digest a wide range of foods. However, the unique signal from food for activating nematodes to digest more diverse food in nature, and what signals activate the animals digestive process are still unclear. Identifying the common digestive signal and its mechanism will help us to understand why the bacterial-feeding nematodes have such great success in nature, which also relates to their essential roles in soil ecosystems.

The nematode *Caenorhabditis elegans* is a free-living roundworm that consumes a diversity of microbes in organic-rich environments such as rotting plant matter (Felix and Braendle, 2010; Samuel et al., 2016; Schulenburg and Felix, 2017), and it is an ideal model for food sensing and food-related behavior research (Shtonda and Avery, 2006). Previously, we established a simple food digestion research system by investigating the effect of food digestion in *C. elegans* on development via feeding the inedible bacteria Staphylococcus saprophyticus (SS) (Geng et al., 2022; Liu and Qi, 2023). *C. elegans* does not grow on either inedible food (*Staphylococcus saprophyticus*) or low-quality food (heat-killed *E. coli*, HK-*E. coli*), but does so when exposed to both (Geng et al., 2022). We have also identified bacterial membrane proteins as signals from low quality food (HK-*E. coli*) as well as the host’s neural and innate immunity pathways that promote digestion of inedible food (*S. saprophyticus*) (Geng et al., 2022). In nature, as *C. elegans* mainly feeds on different species of bacteria, we hypothesized that there may be a common signal from bacteria food that is used as a cue to activate the animal’s food digestive system. This would promote animals to digest inedible food, which can then facilitate the succession of animals in nature by digesting a greater diversity of food.

In this paper, we discovered an unexpected role of bacterial peptidoglycan (PGN) as a food signal in promoting host animals to digest inedible food. Mechanistically, we found that PGN was sensed by conserved intestinal glycosylated protein BCF-1 for inhibiting the mitochondrial unfolded protein response (UPR^mt^), which promotes food digestion. Moreover, activation of UPR^mt^ inhibits food digestion, establishing a connection between the UPR^mt^ and digestion. Together, the data suggest a model in which bacterial PGN acts as a food digestive signal by activating food digestion through a glycosylated protein that regulates intestinal UPR^mt^, facilitating adaptation of the animals under varying food conditions.

## Results

### Low quality food activates animals to digest inedible food, increasing their fitness in nature

Nematodes represent approximately 80% of all the multicellular animals on earth and they have major roles in many ecosystems (Eisenhauer and Guerra, 2019). The free-living nematode *Caenorhabditis elegans* is a major model organism for biological research. In nature, *C. elegans* is particularly abundant in microbe-rich environments, especially rooting plants (Felix and Braendle, 2010; Samuel et al., 2016; Schulenburg and Felix, 2017). These microbes can be used as food for nematodes. If animals could digest a greater diversity of food, they could better survive in their environment.

We first classified food into three classes: good food, inedible food, and low quality food (Figure 1A, a,b,c). We had previously established a simple research system to study food digestion in *C. elegans* (Geng et al., 2022; Liu and Qi, 2023), by observing the growth difference phenotype between in animals fed inedible SS versus HK-*E. coli* with SS together, which makes SS edible (Figure 1A and B). Good food, like laboratory *E. coli-OP50*, can be digested by animals, and will support growth (Figure 1A-a). Inedible food (SS) cannot be digested, and will not support growth (Figure 1A-b, 1B). The low-quality food (HK-*E. coli*), does not support animal growth because it lacks nutrition (Figure 1A-c, 1B), however, the body size on HK-*E. coli* is bigger than on SS feeding, suggesting that animals can digest and use HK-*E. coli* as nutrition (Figure 1B). When low-quality food is mixed with inedible food, the HK-*E. coli* activates the food digestive system of the animals, and they are able to digest SS and grow (Figure 1A-d, 1B). In nature, animals do not always have access to high quality food. As animals could digest more inedible food after eating low quality food, which activates the food digestive system, the animals could have a greater diversity of food to better survive in nature. Our results indicate that there is a mechanism in animals by which low quality food activates its digestive system increasing the fitness of the animals.

**Figure 1.**
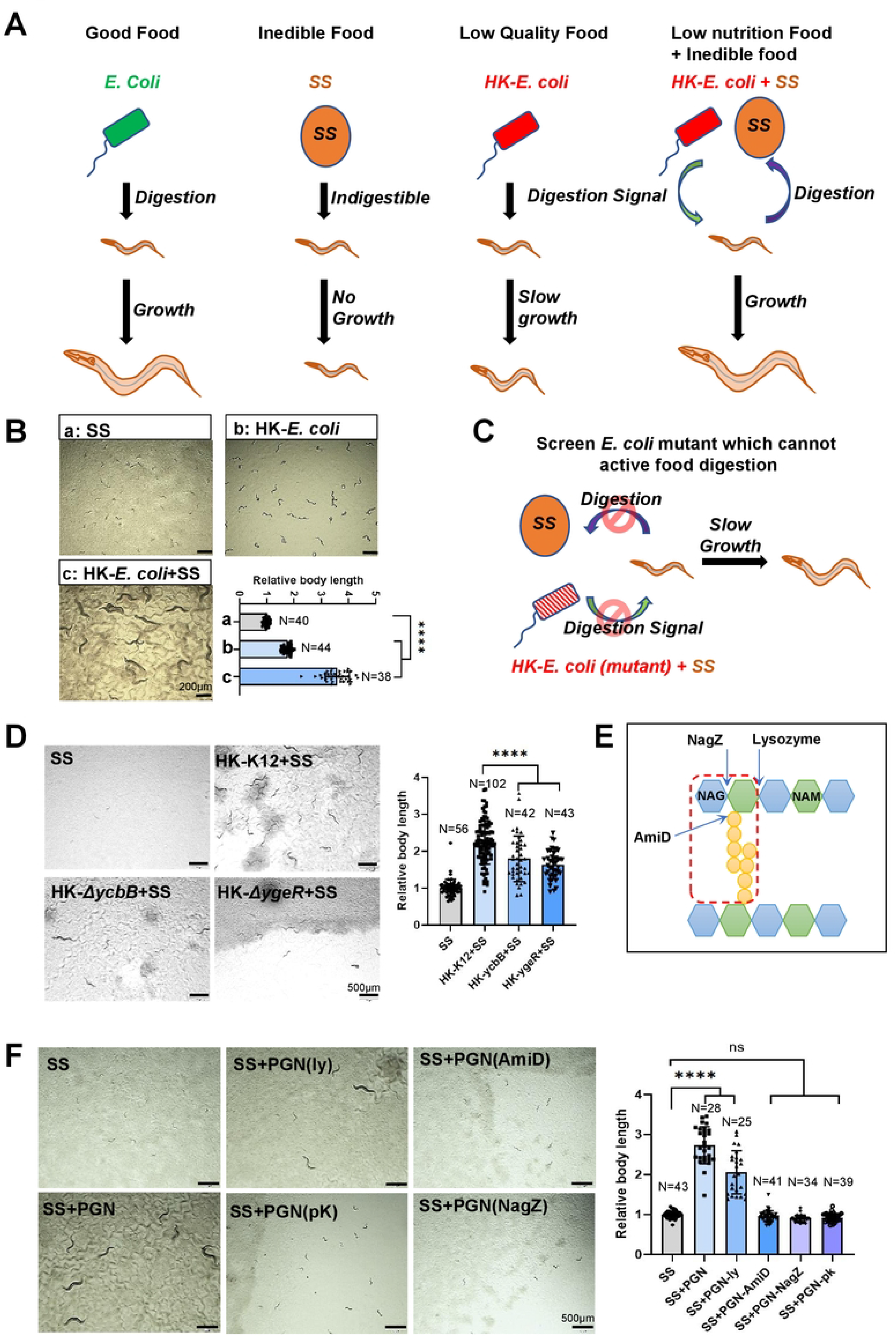
PGN activates *C. elegans* digestive system. (A) Cartoon illustration showing the developmental progression of animals grown on different quality food. Signal from low quality food (HK-*E. Coli)* activates animals to digest inedible food (SS), this mechanism is a potential adaption to increase fitness in animals in nature by increasing the ability to consume a wide range of foods. (B) Developmental phenotype of N2 grown on the SS, HK-*E. Coli*, and HK-*E. Coli*+SS at 20°C for 7d. Scale bar, 500um. (C) Cartoon illustration showing the screening strategy to find the signals from HK-*E. coli* which active animal to digest SS. Synchronized L1 WT animals were grown on HK-*E. coli* (mutant) + SS for 5d to observe the developmental phenotype. *E. coli* mutants lacking the digestive signal would fail to promote animals to digest SS to support growth. (D) Development phenotype of N2 grown on the HK-*E. coli* (*ΔycbB* or *ΔygeR* mutant) + SS or SS at 20°C for 5d post-L1 synchronization. Scale bar, 500um. (E) Schematic representation of the peptidoglycan structure and cleavage points of enzymes by arrows. Red box indicates the structure for the digestion signal from PGN which contains 5’NAG-NAM disaccharide muropeptides with an amino acid peptide attached to NAM. NagZ, glucosaminidase; AmiD, amidase; NAG, 3N’-acetylglucosamine; NAM, 5N’-acetylmuramic acid. (F) Developmental progression of animals grown on SS+ enzyme-treated PGN at 7d at 20°C. Scale bar, 500um. Developmental progression of animals is scored by relative worm body length. For all panels, N= number of animals which were scored from at least three independent experiments. Data are represented as mean ± SD. ****p<0.0001, *** P < 0.001, **P < 0.01, *P < 0.05, ns: no significant difference.

### Bacterial peptidoglycan activates animals to digest SS food

We have shown that HK-*E. coli* activates animals to digest SS (Geng et al., 2022). We next screened for *E. coli* mutants that would fail to promote animals to digest SS to support growth (Figure 1C). After screening, we found that worm growth was significantly slower when feeding HK-*E. coli* mutants (ycbB and ygeR)+SS (Figure 1D), indicating that the ability to digest food is decreased in animals when individually fed these HK-mutant *E.coli*. YcbB has L,D-transpeptidase activity for peptidoglycan (PGN) maturation, and YgeR is putatively involved in PGN hydrolysis (Tian and Han, 2022). Therefore, we asked whether *E. coli* PGN itself activates digestion to promote worms to digest SS, by adding extracted PGN to SS. We found that the slow growth phenotype of worms fed SS was suppressed (Figure 1F), suggesting that PGN activates food digestion in worms.

PGN, the core and unique component of all bacterial cell walls, is a polymer of β(1–4)-linked N-acetylglucosamine (GlcNAc) and N-acetylmuramic acid (MurNAc), crosslinked by short peptides. To identify the specific structural unit of PGN which activates food digestion, we employed several enzymes which cleave PGN at specific sites (Figure 1E). We found that PGN treated with N-acetylmuramoyl-L-alanine amidase AmiD, Beta-hexosaminidase NagZ, (Figure S1) or Protease K could no longer activate the animals to consume SS (Figure 1F). However, lysozyme treated PGN still promoted animals to consume SS for supporting growth (Figure 1F). These results suggest that the specific PGN molecules effective in activating food digestion are 5’NAG-NAM disaccharide muropeptides with an amino acid peptide attached to NAM.

### Screen for PGN-binding proteins involved in food digestion

In order to determine how PGN activates food digestion, we began by investigating whether *E. coli-*binding proteins (He et al., 2023) and PGN-binding proteins (Tian and Han, 2022) previously implicated in food digestion have a role. 44 proteins were identified as both *E. coli*-binding and PGN-binding (Figure 2A). Among these 44 genes, 23 genes are expressed in intestine, which is the main digestive organ in animals (Figure 2A). If the particular PGN-interaction protein is critically involved in food digestion, then RNAi knockdown of this candidate may generate a slow growth phenotype in animals fed HK-*E. coli*+SS. We therefore performed an RNAi screen of all 23 candidate genes (Figure S2) and found that animals fed with *bcf-1* RNAi grew slowly on HK-*E. coli*+SS (Figure 2B). Furthermore, two independent *bcf-1* mutants also grew slowly when fed HK-*E. coli*+SS (Figure 2C). These data suggested that potential PGN-binding protein, BCF-1, is involved in promoting digestion of SS food in animals.

**Figure 2.**
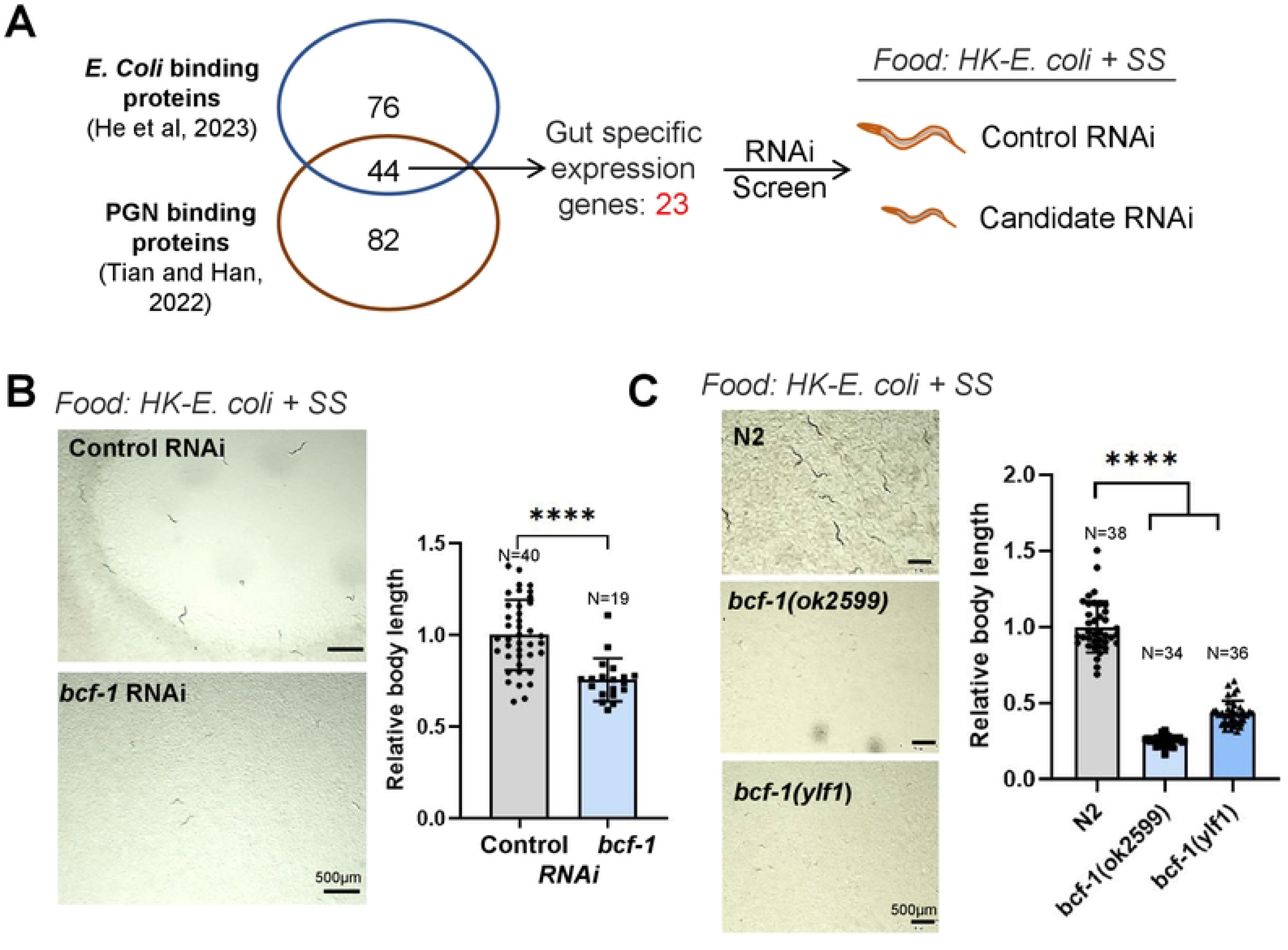
Screening PGN-Binding proteins involved in food digestion. (A) Cartoon illustration showing the screening strategy to find the PGN-binding protein which active animals to digest SS. Venn diagram indicating the total number of identified *E. coli* binding proteins (He et al., 2023) and PGN binding proteins (Tian and Han, 2022) and their overlap. From 44 overlap genes, 23 intestinal specific genes were tested by RNAi screening. RNAi knockdown of the candidate is expected to generate a slow growth phenotype in animals fed HK-*E. coli*+SS. (B-C) Developmental progression of animals with *bcf-1* RNAi (B) or *bcf-1* mutations (C) grown on HK-*E. coli* +SS. Developmental progression of animals is scored by relative worm body length. For all panels, N= number of animals which were scored from at least three independent experiments. Data are represented as mean ± SD. ****p<0.0001, *** P < 0.001, **P < 0.01, *P < 0.05, ns: no significant difference.

### PGN interacts with BCF-1 for food digestion

Our previous finding showed that BCF-1 is an N-glycosylated intestinal protein, which regulates *E. coli* colonization by directly binding bacteria (He et al., 2023). Also, in the HK-*E. coli*+SS feeding condition, BCF-1 is only expressed in the intestine (Figure 3A). Animals with intestinal *bcf-1* RNAi grew slowly on HK-*E. coli*+SS (Figure 3B), suggesting that intestinal BCF-1 activates food digestion.

**Figure 3.**
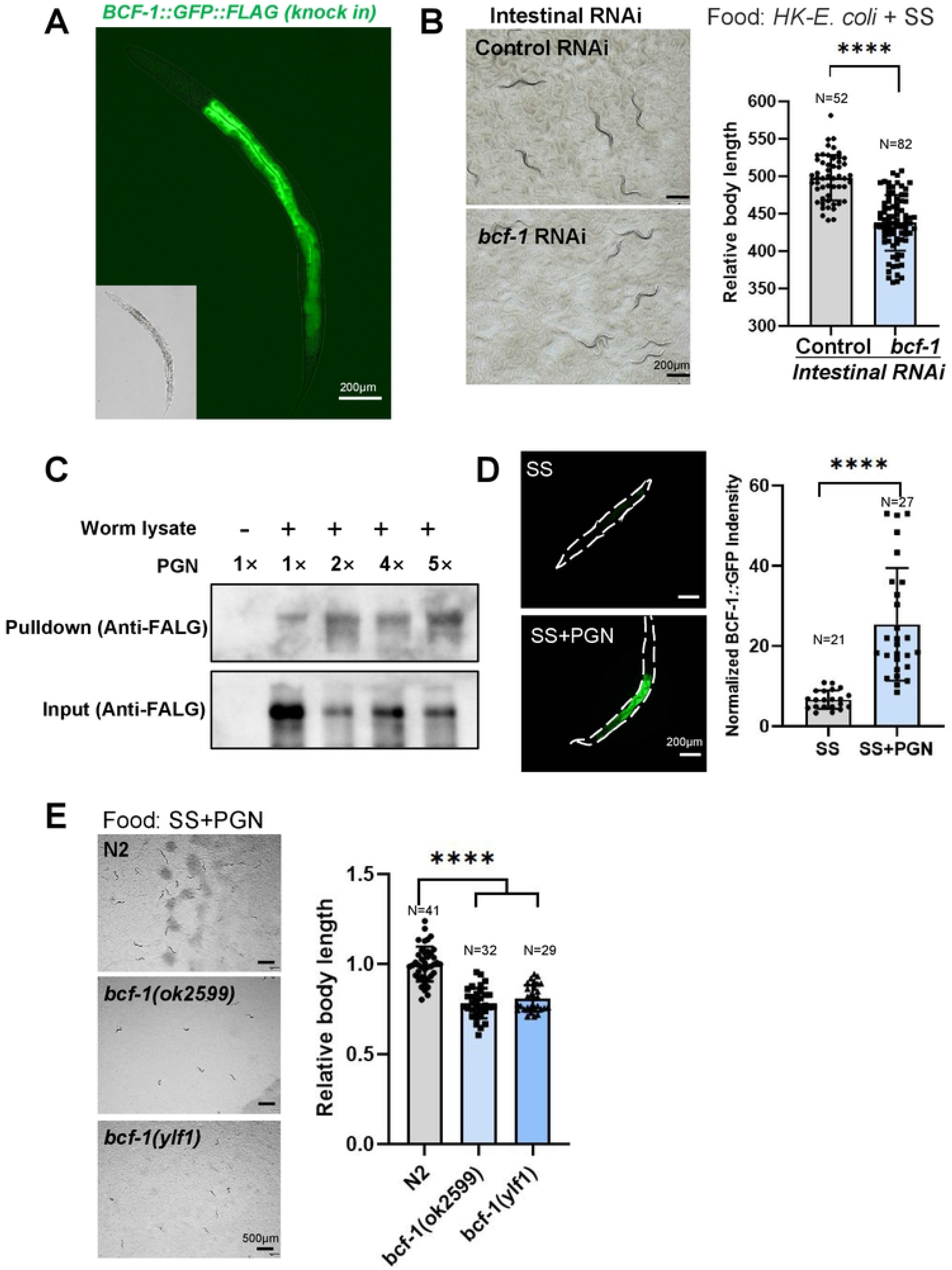
PGN activates food digestion through BCF-1. (A) Fluorescence image showing that BCF-1 specifically expressed in the intestine by using a single-copy insertion of the *bcf-1p::bcf-1::gfp::flag* reporter strain. (B) Developmental progression of animals with intestinal specific RNAi *bcf-1* grown on HK-*E. coli* +SS at 4d at 20°C. (C) *In vitro* PGN binding assay (pull-down assay) showing that PGN interacts with BCF-1 protein. BCF-1 from worm lysates (*bcf-1p::bcf-1::gfp::flag* reporter strain) was bound to PGN and the binding increased in a concentration-dependent manner. Anti-Flag was used as an input control, which indicate that PGN was incubated with an all most equivalent amount of BCF-1 tagged (BCF-1-GFP-FLAG) proteins. (D) Microscope images and bar graph showing that BCF-1::GFP expression is induced in the *bcf-1p::bcf-1::gfp::flag* reporter strain when fed with SS+PGN. (E) Developmental progression of the *bcf-1* mutant grown on SS+PGN at 5d at 20°C. Developmental progression of animals is scored by relative worm body length. For all panels, N= number of animals which were scored from at least three independent experiments. Data are represented as mean ± SD. ****p<0.0001, *** P < 0.001, **P < 0.01, *P < 0.05, ns: no significant difference.

We then asked if PGN-activated food digestion is dependent on its interaction with BCF-1 protein. First, we incubated PGN with worm lysate (BCF-1::FLAG) and found that PGN bound to BCF-1 in a concentration-depended manner (Figure 3C), suggesting that PGN interacts with BCF-1. Secondly, we found that BCF-1 expression is induced by PGN in the SS food feeding condition (Figure 3D), suggesting that PGN activates food digestion through promoting BCF-1 expression. Third, *bcf-1* mutants grew slowly on SS+PGN (Figure 3E), suggesting that PGN promotes SS digestion which requires BCF-1. Together, these results indicated that PGN activates food digestion dependent on its interaction protein BCF-1 in the intestine. As well, *C. elegans* BCF-1 is a conserved glycosylated protein found only in bacterial-feeding nematodes (Figure S3) including in *C. remanei, C. nigoni, C. briggsae, C. bovis*, *and C. auriculariae*. Thus, our findings indicated that conserved nematode BCF-1 could sense unique bacterial PGN to activate food digestion in animals, which facilitate adaptation of the nematodes in their natural environment by increasing their ability to consume a wide range of bacterial foods.

### PGN inhibits UPR^mt^ through BCF-1 for food digestion

Bacterial peptidoglycan muropeptides benefit mitochondrial homeostasis by acting as ATP synthase agonists in the inhibition of the mitochondrial unfolded protein response (UPR^mt^) (Tian and Han, 2022). We found that UPR^mt^ was induced in worms fed with the PGN mutant*ΔycbB* (Figure 4A). We thus wondered if PGN promotes food digestion through inhibition of UPR^mt^ by acting with BCF-1. If the PGN-binding protein BCF-1 is involved in the beneficial impact of PGN on mitochondrial homeostasis, then the *bcf-1* mutation may generate phenotypes similar to that caused by the PGN mutant diet, which induces UPR^mt^ and results in increased food avoidance behavior (Tian and Han, 2022). As we hypothesized, UPR^mt^ is induced in animals with *bcf-1* RNAi (Figure 4B) or a *bcf-1* mutation (Figure 4A), and food avoidance behavior is also increased in the *bcf-1* mutant (Figure 4C), indicating that mutation of BCF-1 shows similar phenotypes caused by PGN deficiency.

**Figure 4.**
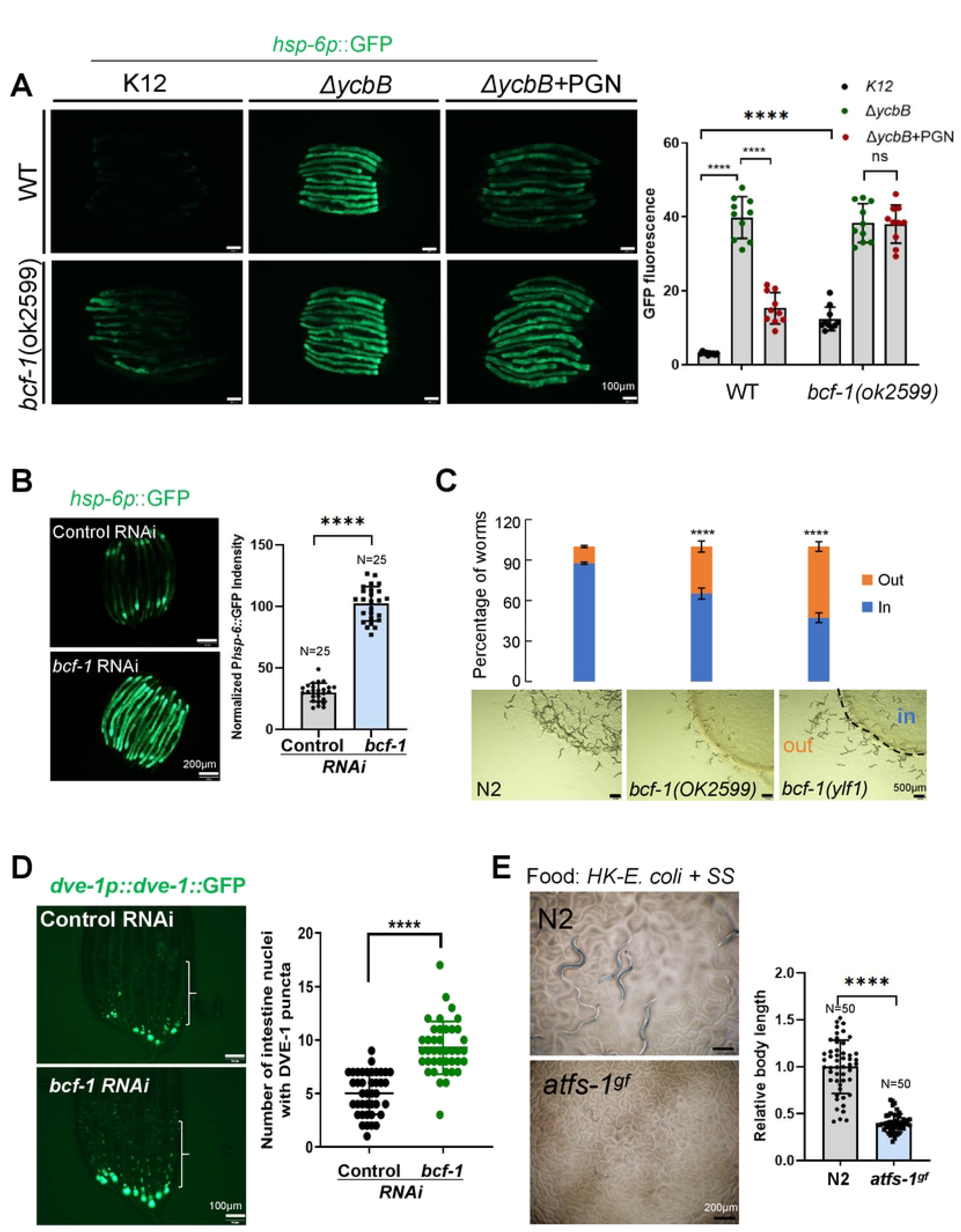
PGN inhibits UPR^mt^ through *bcf-1* for food digestion. (A) Microscope image and bar graph showing the role of PGN in inhibiting UPR^mt^ is dependent on BCF-1. UPR^mt^ was activated by feeding the PGN mutant (*ΔycbB*), and was suppressed by the addition of PGN. However, the addition of PGN failed to suppress UPR^mt^ in the *bcf-1* mutant. (B) Fluorescence image and bar graph showing that UPR^mt^ was induced in animals with *bcf-1* RNAi treatment. (C) The food avoidance phenotype of *bcf-1* mutants. Food avoidance is increased in animals with *bcf-1* mutation. (D) Fluorescence image and bar graph showing that DVE-1::GFP was accumulated in intestinal nuclei in *dve-1p::dve-1::gfp* animals with *bcf-1* RNAi treatment. (E) Developmental progression of *atfs-1(gf)* grown on HK-*E. coli* +SS. For all panels, N= number of animals which were scored from at least three independent experiments. Data are represented as mean ± SD. ****p<0.0001, *** P < 0.001, **P < 0.01, *P < 0.05, ns: no significant difference.

Next, we asked if PGN’s function in inhibition of UPR^mt^ through BCF-1. We found that adding PGN to the animals fed with*ΔycbB* could suppress the UPR^mt^, and that this suppression effect was eliminated when the PGN was added to *bcf-1* mutant worms (Figure 4A), suggesting that PGN-BCF-1 interaction plays a critical role in maintaining the UPR^mt^.

Finally, we wondered if a balanced UPR^mt^ is critical for food digestion. We found that DVE-1, a critical cofactor in ATFS-1-mediated UPR^mt^ is translocated into the nucleus in the animals with *bcf-1* RNAi (Figure 4D) which also show a decreased ability to digest food (Figure 2B-C). Thus, activation of UPR^mt^ could decrease the ability to digest food in animals. Activation of ATFS-1, the key factor in the UPR^mt^ pathway, is sufficient to activate UPR^mt^. We found that the *afts-1*(*et18, gain of function*) mutant, which has constitutive activation of UPR^mt^, grew significantly slower on HK-*E. coli* + SS (Figure 4E), suggesting that activation of UPR^mt^ inhibits food digestion. Together, these data revealed a mechanism by which PGN-BCF-1 interaction plays a critical role in activating food digestion through UPR^mt^ inhibition.

## Discussion

The succession of bacterial-feeding nematode species in nature plays an important role in ecosystems (Eisenhauer and Guerra, 2019). Thus, the ability to digest a wide range of foods plays a pivotal role for survival. However, common signals from the bacterial food which activates host food digestion are still unclear. In this study, we re-evaluated our previous finding that low quality food (HK-*E. coli*) promotes animals to digest inedible food (SS) (Geng et al., 2022), which may increase the ability of animals to consume a greater variety of food. This fascinating result may underlie a mechanism for increased fitness of nematodes in nature. By using this established digestion system, and screening of bacterial mutants, we found that bacterial peptidoglycan stimulates animals to digest an inedible food, SS. On the host side, we found that intestinal glycosylated protein, BCF-1, interacts with PGN to promote food digestion via inhibiting UPR^mt^. As well, UPR^mt^ was activated in animals by feeding mutant bacteria lacking PGN or by *bcf-1* mutation, both which showed defects in food digestion. We also found that constitutive activation of UPR^mt^ is sufficient to inhibit food digestion, which established the connection between the UPR^mt^ and digestion. Therefore, our study uncovered a fascinating mechanism in bacterial-feeding nematodes for surviving in nature by digesting more complex food via sensing bacterial PGN through glycosylated protein in the intestine (BCF-1) which inhibits UPR^mt^ in host (Figure 5). This study also suggests the beneficial roles of bacterial PGN in activating the food digestive system, as the abundant amount of commensal gut bacteria which contain the PGN in their cell wall in the animal kingdom.

**Figure 5.**
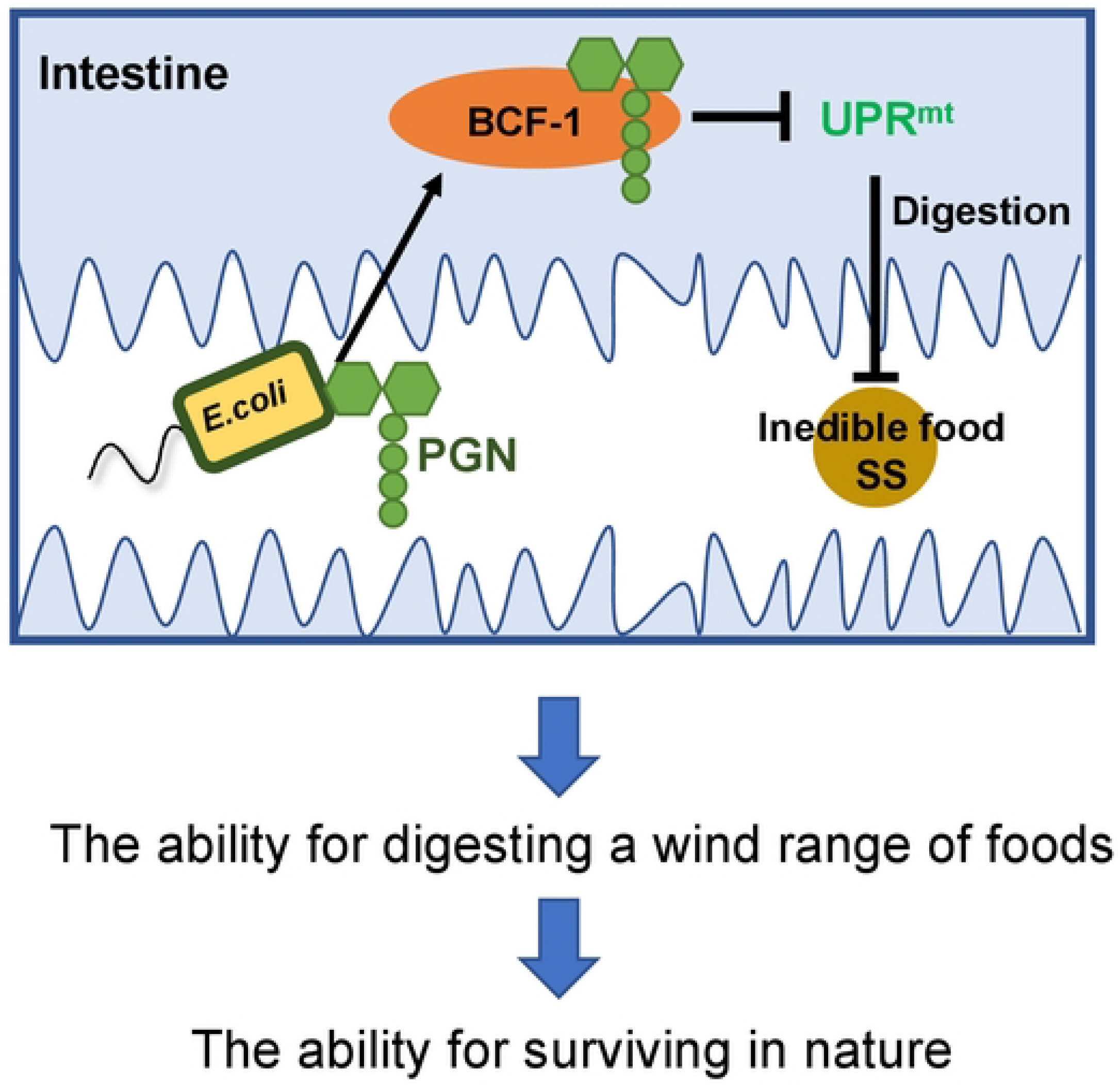
Survival strategy in animals by sensing unique bacterial PGN for activating food digestion. A model showing that conserved glycosylated protein (BCF-1) in nematode interacts with bacterial PGN to promote digest inedible food (SS) via inhibiting UPR^mt^. This is a fascinating mechanism in animals for surviving in nature by increasing the ability to digest a wide range of foods through sensing the unique bacterial cell wall component PGN.

Nematodes are ubiquitous organisms that have a significant global impact on ecosystems, agriculture, and human health (Murfin et al., 2012). Bacterial-feeding nematodes are primarily responsible for being beneficial in decomposition and nutrient cycling in ecological and agricultural systems (Freckman, 1988). Thus, consumption and digestion of a wide range of foods/bacteria are essential for nematodes to maintain physiological integrity and live in a healthy state for surviving in nature. The microbes as food for nematodes are composed of both gram-positive (G+) and gram-negative (G-) bacteria. The cell walls of bacteria contain many components that are recognized by the animals for regulating innate immune responses, such as lipopolysaccharide (Hajjar et al., 2002; Shi et al., 2014; Wright et al., 1990), ADP-heptose (Zhou et al., 2018), and peptidoglycan (PGN) (Gupta, 2008; Wolf et al., 2016). PGN is a core and unique component of all bacterial cell walls. Evolutionarily, bacterial-feeding nematodes could evolute a mechanism to sense bacterial PGN as component of food for maintaining its state of health, which is a survival mechanism for animals living in nature. Here, we found that the food digestive system was activated by a unique microbe-associated pattern, PGN, in the nematode *Caenorhabditis elegans*, promoting animals to consume more complex food including normally inedible food. We also show that conserved glycosylated protein (BCF-1) in nematodes (including *C. remanei, C. nigoni, C. briggsae, C. bovis, C. auriculariae*, *and C. elegnas*), interacts with PGN to promote food digestion via inhibiting the mitochondrial unfolded protein response (UPR^mt^), which is a conserved transcriptional response activated by multiple forms of mitochondrial dysfunction. Therefore, the conserved BCF-1 in nematodes could sense a bacterial unique PGN to activate food digestion through regulating UPR^mt^ in the host. This mechanism of food digestion has only been seen in bacterial-feeding nematodes as PGN binding protein BCF-1 does not have homology in other species, including humans. This is the efficient survival rule of eating more diverse food and having a higher fitness for bacterial-feeding nematodes as these animals use bacterial as food. Thus, the beneficial role of PGN for nematodes food digestion through BCF-1 could be a causal mechanism for co-evolution between microbe and nematodes in adaptation of food diversity. Recently studies have found that bacterial peptidoglycan muropeptides benefit mitochondrial homeostasis and animal physiology by acting as ATP synthase agonists in *C. elegans* (Tian and Han, 2022) and that peptidoglycan components (MurNAc-L-Ala and MurNAc) also enhance pathogen tolerance in *C. elegans* (Rangan et al., 2016) which points to the beneficial role of PGN in the animals. Therefore, these studies (Rangan et al., 2016; Tian and Han, 2022) also support our hypothesis that co-evolution between microbial PGN and nematodes in adaptation in the natural environment including diversity food and pathogens.

Collectively, this study reveals that unique bacterial PGN was sensed by conserved glycosylated protein (BCF-1) for food digestive system activation in nematodes, which is important for animals in nature enabling them to digest more diverse food, increasing their fitness. In addition, by sensing PGN the food digestive mechanism reported here could have a significant influence on our understanding of the mechanism of how animals adapt to the abundance of food diversity by regulating its digestive system and feeding behaviors.

## Author Contributions

F. H and H. L performed most experiments and analyzed data. B.Q. designed research, supervised this study, and wrote the paper with inputs from F. H and H. L.

## Acknowledgments

We thank the Caenorhabditis Genetics Center (CGC) (funded by NIH P40OD010440) for strains; Dr. Leonard Krall for editing services; Dr. Zhao Shan for suggestions. This work was supported by the Ministry of Science and Technology of the People’s Republic of China (2019YFA0803100, 2019YFA0802100), the National Natural Science Foundation of China (32170794), Yunnan Applied Basic Research Projects (2019FY003022, 202001AV070011, 202001AW070006, 202201AT070196), the Yunnan University Startup Program.

## Supplemental Figure legends

**Figure S1.** SDS-PAGE gel showing purified recombinant proteins (AmiD and NagZ). Related to Figure 1.

**Figure S2. RNAi screen of gut specific expression genes which may be involved in digestion. Related to Figure 2.**

(A-B) Developmental phenotype (A) and quantification of body length (B) of animals with indicated RNAi grown on HK-*E. coli*+SS for 4 days at 20°C.

**Figure S3. Phylogenetic tree of BCF-1 in nematodes. Related to Figure 3.** Phylogenetic tree showing that *C. elegans* BCF-1 protein is conserved in nematodes, especially in *C. remanei, C. nigoni, C. briggsae, C. bovis, C. auriculariae*, *and C. angaria*.

## Star Methods

### Resource Availability

#### Lead contact

Further information and requests for reagents may be directed to the Lead contact Bin Qi.

#### Materials availability

All reagents and strains generated by this study are available through request to the lead contact with a completed Material Transfer Agreement.

#### Data and code availability

Original data in this paper has been deposited in Mendeley Data (doi: 10.17632/zhtwwb7s3z.1).

This paper does not report original code.

Any additional information required to reanalyze the data reported in this paper is available from the lead contact upon request.

### Experimental model and subject details

#### *C. elegans* strains and maintenance

Nematode stocks were maintained on nematode growth medium (NGM) plates seeded with bacteria (*E. coli* OP50) at 20°C.

The following strains/alleles were obtained from the Caenorhabditis Genetics Center (CGC) or as indicated:

N2 Bristol (wild-type control strain);

RB1971[*F57F4.4(ok2599)];*

*SJ4100: zcls13 [hsp-6p::GFP + lin-15(+)];*

*MGH171: sid-1(qt9) V, alxIs9 [vha-6p::sid-1::SL2::GFP]*

*SJ4197: zcIs39 [dve-1p::dve-1::GFP]*.

*QC118: atfs-1(et18)*

YNU30: *bcf-1(ylf1)*;

PHX4067: *syb4067[bcf-1p::bcf-1::3xflag::gfp(knock-in)*];

*bcf-1(ok2599), hsp-6p::GFP was constructed by crossing*

RB1971[*F57F4.4(ok2599)] with SJ4100: zcls13 [hsp-6p::GFP + lin-15(+)]*.

#### Bacterial strains

*E. coli-*OP50, *E. coli-K12* (BW25113), *E. coli-K12* mutant, and *Staphylococcus saprophyticus* were cultured at 37°C in LB medium. A standard overnight cultured bacteria was then spread onto each Nematode growth media (NGM) plate.

## Method Details

### Heat-killed *E. Coli* preparation

Heat-killed *E. coli* was prepared by an established protocol (Qi et al., 2017). A standard overnight bacterial culture was concentrated to 1/10 vol and was then heat-killed at 80°C for 2h.

### Preparation of *S. saprophyticus* (ATCC 15305) and HK-E. *coli*+S. saprophyticus (SS)

*S. saprophyticus* and HK-*E. coli*+SS were prepared by following our established protocol (Geng et al., 2022). For SS preparation, a standard overnight culture of SS (37°C in LB broth) was diluted into fresh LB broth (1:100 ratio). SS was then spread onto each NGM plate when the diluted bacteria grew to OD600= 0.5. For HK-*E. coli*+SS preparation, HK-*E. coli* (50ul) and *SS* (50ul) were mixed at a 1:1 ratio, then 100ul of the mixture was spread onto NGM plates.

About 100-200 synchronized L1 worms were then seeded onto the indicated plates (SS, or HK-*E. coli*+SS) and cultured at 20 °C for growth phenotype obversion.

### *E. coli* Keio collection screen

*E. coli* mutants are from the Keio *E. coli* single mutant collection (Baba et al., 2006). Mutant bacteria strains, as well as the wild-type control strain BW25113, were cultured overnight in LB medium with 50 ug/ml kanamycin in 96-well plates at 37°C. Overnight cultured bacteria were then heat-killed following our established protocol (Qi et al., 2017). *S.saprophyticus* was also prepared as described above. HK-*E. coli* mutants (50ul) and *S.saprophyticus* (50ul) were mixed at a 1:1 ratio and then 100 ul of the mixture was spread onto 3.5 cm NGM plates. About 100-200 synchronized L1 worms were then seeded onto HK-*E. coli*+SS feeding plates and cultured for 3-4 days at 20 °C. The animals size was then observed.

### RNAi treatment

All feeding RNAi experiments used bacterial clones from the MRC RNAi library (Kamath et al., 2003) or the ORF-RNAi Library (Rual et al., 2004). RNAi plates were prepared by adding IPTG with final concentration of 1mM to NGM. Overnight cultured RNAi strains (LB broth containing 100 ug/ml ampicillin) and the control strain (HT115 strain with empty L4440 vector) were seeded into RNAi feeding plates and cultured at room temperature for 2 days before use.

### RNAi screen

Synchronized L1 worms were treated by feeding candidate RNAi for the second generation and grew to adult. They were then bleached and allowed to hatch in M9 buffer for 18hr. The synchronized L1 worms were seeded on the feeding plate (HK-*E. coli*+SS). The worm development phenotype (body length) was measured after culturing 3-4 days at 20 °C.

### Food avoidance behavior assay

Food avoidance assay was performed according to published method (Tian and Han, 2022). 20ul of overnight cultured *E. coli-OP50* was seeded on the center of 6cm NGM plates. About 100 synchronized L1 animals were seeded onto the bacterial lawns and cultured at 20°C for 48 hours. The avoidance index was determined by N_off_/_on_/N_total_.

### PGN extraction, enzyme treatment and supplementation assays

PGN was extracted according to an established method (Tian and Han, 2022). In brief, bacterial pellets from a standard overnight culture of *E. Coli* OP50 (37°C in LB broth) were resuspended in 1/10 vol 1M NaCl solution and boiled at 100°C in a heating block for 30min. After washing 3 times by ddH_2_O, the insoluble cell fragments were crushed by ultrasound for 1h, followed by stepwise digestion using DNAse (50ug/ml), RNAse (60ug/ml), and trypsin (50ug/ml) at 37°C for 1h, respectively. After collecting bacterial pellets and washing them with ddH_2_O three times, they were heated at 100 °C for 5 min to inactivate the enzymes. Finally, the pellets collected by centrifugation at 12,000g was re-suspended with HEPES buffer (20mM, pH=7.5) and stored at -20°C.

For enzymatic treatment of isolated PGN, lysozyme (500ug/ml), NagZ (purified from BL21 *E. coli*,500ug/ml), AmiD (purified from BL21 *E. coli*, 500ug/ml), or Protease K (500ug/ml) were added to isolated PGN and incubated at 37°C for 24h. The reaction was terminated by adding trypsin (500ug/ml) at 37°C for 2h. Finally, the above samples were heated at 100°C for 5 min to inactivate the enzymes.

For supplementation assays, indicated bacteria (SS, *ΔycbB*) and PGN solution mixtures (1:1 ratio) were seeded on NGM plates. The synchronized L1 worms were then seeded on the indicated plate (SS, SS+PGN, or SS+enzyme treated PGN; *ΔycbB, ΔycbB*+PGN).

### Protein expression and purification

The expression of recombinant proteins (NagZ and Amid) was performed by established protocols (Tian and Han, 2022). Briefly, the NagZ and AmiD genes were amplified by PCR from *E. coli-K12* (BW25113) genomic DNA. Then the target genes were inserted into pET28a vector by homologous recombination. These constructs were transformed into *E. coli BL2*1(DE3). Overnight cultured BL21 bacteria was diluted into fresh LB broth (1/100 vol ratio) and cultured to OD600≈ 0.6. Then, the expression of recombinant proteins was induced by 1 mM IPTG at 20°C for 19 h. Bacterial pellets were collected by centrifugation at 5,000g for 10min at 4°C, and the pellets were lysed in buffer (25mM Tris-HCl pH=8.0, 150Mm NaCl, 10% Glycerol, 0.1% NP40). The suspension was ultrasonically broken (25%power, followed by centrifugation at 12,000g for 30min at 4°C. The supernatants were incubated with Ni-NTA agarose beads (TianGen 30210) and rinsed 3 times with a washing buffer (25mM Tris-HCl pH=8.0, 150mM NaCl, 10% Glycerol, 0.1% NP40, 30mM Imidazole). Finally, the Elution buffer (25mM Tris-HCl pH=8.0, 150mM NaCl, 10% Glycerol, 0.1% NP40, 500mM Imidazole) was used to eluted the recombinant protein. Amicon Ultra centrifugal filters (10kD) were used to concentrate the proteins and exchange buffer (25 mM HEPES pH 7.5, 250 mM NaCl, and 1 mM DTT).These proteins were stored at -80°C.

### Worm total protein extraction

Total proteins were extracted according to our established method (He et al., 2023). Worms were lysed by freeze-thawing three times in protein lysis buffer containing PMSF (1mg/ml) in liquid nitrogen. The worms were then ground in a tissue grinding tube. The resultant liquid was then centrifuged at 12,000rpm for 30min at 4°C, and the supernatant was taken as the total protein of worms. The protein concentration was measured by using the Pierce BCA protein assay kit (ThermoFisher, 23227).

### BCF-1 expression assay with PGN induction

Synchronized L1 animals (*bcf-1p::bcf-1::gfp::flag*) were seeded onto SS or SS+PGN feeding plates and cultured at 20 °C 4 days before observing the fluorescence.

### PGN-protein binding assay

The PGN-protein binding assay was carried out as described (Cash et al., 2006). Briefly, total protein (1mg) of *C. elegans* was incubated with PGN at 4°C for 4h. At the end of incubation, the PGN was pelleted, then the pellet was washed 3 times with washing buffer (100mM NaCl, 50mM Tris-HCl pH=7.5). Proteins associated with the pellet and the incubated supernatant were detected by Western blot with an anti-flag antibody.

### Analysis of the fluorescence intensity in worms

For fluorescence imaging (*hsp-6p::gfp*), worms were anesthetized with 25 mM levamisole and photographs were taken using an Olympus MVX10 dissecting microscope with a DP80 camera. To quantify fluorescent intensity, the entire intestine regions were outlined and quantified using ImageJ software. The fluorescence intensity was then normalized to body area.

For quantifying BCF-1::GFP, the worms were mounted on 2% agarose pads with 25 mM levamisole and imaged using an Olympus MVX10 microscope with a DP80 camera.

## Quantification and Statistical Analysis

### Quantification

ImageJ software was used for quantifying fluorescence intensity of various of reporters or analysis of developmental body length of nematodes. The fluorescence intensity was normalized to body area.

### Statistical analysis

All statistical analyses were performed using Student’s t-test. Two-tailed unpaired t test was used for statistical analysis of two groups of samples. Data are presented as Mean ± SD, and p<0.05 was considered a significant difference. For all figures, ‘‘n’’ represents number of animals which were scored from at least three independent experiments.

